# *Botrytis cinerea* small RNAs are associated with tomato AGO1 and silence tomato defense-related target genes supporting cross-kingdom RNAi

**DOI:** 10.1101/2022.12.30.522274

**Authors:** Baoye He, Qiang Cai, Arne Weiberg, Wei Li, An-Po Cheng, Shouqiang Ouyang, Katherine Borkovich, Jason Stajich, Cei Abreu-Goodger, Hailing Jin

## Abstract

Cross-kingdom or cross-species RNA interference (RNAi) is broadly present in many interacting systems between microbes/parasites and their plant and animal hosts. A recent study by Qin *et al*. (2022) performed correlation analysis using global sRNA- and mRNA-deep sequencing data of cultured *B. cinerea* and *B. cinerea*-infected tomato leaves and claimed that cross-kingdom RNAi may not occur in *B. cinerea*–tomato interaction (Qin *et al*., 2022). Here, we use experimental evidence and additional bioinformatics analysis of the datasets produced by Qin *et al*. (2022) to identify the key reasons why a discrepancy between the conclusion of Qin et al. 2022 and previously published findings occurred. We also provided additional experimental evidence to support the presence of cross-kingdom RNAi between tomato and *B. cinerea*. We believe it is important to clarify the basic concept and mechanism of cross-kingdom/cross-species sRNA trafficking and illustrate proper bioinformatics analyses in this regard for all the scientists and researchers in this field.

## 1 INTRODUCTION

Cross-kingdom or cross-species RNA interference (RNAi) is a phenomenon in which small RNAs (sRNAs) are transported between interacting organisms to silence the expression of target genes in the adversary or interacting partner (Weiberg *et al*., 2013, Cai *et al*., 2018, Weiberg *et al*., 2014). Our lab first discovered this communication mechanism between plants and the fungal pathogen *Botrytis cinerea*. Some sRNAs of *B. cinerea* are transported into host plants where they hijack host RNAi machinery for cross-kingdom gene silencing (Weiberg et al., 2013). We found that the transferred fungal sRNAs are loaded into the host Arabidopsis Argonaute 1 (AGO1) protein to silence host target genes involved in immune responses (Weiberg et al., 2013). This phenomenon has since been observed later in many different interacting organisms. For example, other plant fungal pathogens (such as *Fusarium oxysporum* and *Verticillium dahliae*), oomycete pathogens (such as *Hyaloperonospora arabidopsidis*), the parasitic plant *Cuscuta campestris*, and even the symbiotic ectomycorrhizal fungus *Pisolithus microcarpus* and the bacterial symbiont Rhizobium (*Bradyrhizobium japonicum*) can send a set of sRNAs into the host plant to silence the expression of host target genes (Dunker *et al*., 2020, Shahid *et al*., 2018, Ren *et al*., 2019, Wong-Bajracharya *et al*., 2022, Ji *et al*., 2021, Zhang *et al*., 2022). Furthermore, the mode of action of transferred microbial sRNAs seems to be conserved. The sRNAs from *F. oxysporum* (Ji et al., 2021), *V. dahliae* (Wang et al., 2016), *H. arabidopsidis* (Dunker et al., 2020) and the *Rhizobium* (Ren et al., 2019) also utilize host AGO proteins for silencing host target genes. Excitingly, cross-kingdom or cross-species RNAi has also been observed in animal-pathogen or parasite interaction systems. For example, the fungal pathogen of mosquito, *Beauveria bassiana* transfers a microRNA-like RNA (milRNA) to the host cells. This milRNA is loaded into mosquito AGO1 to silence the host immunity gene *Toll receptor ligand Spätzle 4* (Cui *et al*., 2019).

Some parasites, such as the gastrointestinal nematode *Heligmosomoides polygyrus*, send a range of its sRNAs into mammalian gut cells to target and silence immunity and inflammation-related genes (Buck *et al*., 2014, Chow *et al*., 2019). The causal bacterial pathogen for human Legionnaires’ disease, *Legionella pneumophila* also sends bacterial sRNAs into human host cells to regulate the expression of innate immune response genes (Sahr *et al*., 2022). Thus, cross-kingdom/cross-species RNAi is widely observed and accepted by scientists studying many interaction systems between microbes/pests and their plant or animal hosts.

Recently, Qin *et al*. (2022) performed sRNA- and mRNA-deep sequencing analysis of cultured *B. cinerea* and *B. cinerea*-infected tomato leaves and conducted inoculation assays with mutated *B. cinerea* strains that deleted a transposon or two Dicer-like genes (*bcdcl1dcl2*). They claimed that cross-kingdom RNAi does not occur in *B. cinerea*–tomato interaction (Qin *et al*., 2022). Here, we re-analyzed RNA sequencing data released by this study and aimed to recapitulate infection assay results to identify the key reasons for the discrepancy with previously published findings. We also provided additional experimental evidence that *B. cinerea* sRNAs are associated with tomato AGO1 to silence tomato target genes, which supports the presence of cross-kingdom RNAi between tomato and *B. cinerea*. We believe it is important to clarify the basic concept and mechanism of cross-kingdom/cross-species RNAi and to illustrate the proper bioinformatics analyses of sRNA and mRNA deep sequencing libraries.

## 2 RESULTS

### 2.1 Discrepancies in the bioinformatics analysis and interpretation of sRNA and mRNA sequencing data

Qin *et al*. (2022) performed a correlative analysis on sRNA and mRNA transcriptome datasets of cultured *B. cinerea* and *B. cinerea-infected* tomatoes. They identified a total of 27,918 unique *B. cinerea* sRNAs with sizes of 20-24 nt from *B. cinerea*-infected tomato leaves harvested at 12, 16, and 24 h post-infection (hpi) as well as from *B. cinerea* grown in liquid culture. They used the entire 27,918 sRNA set, 70-90% of which derived from *B. cinerea* ribosomal RNAs (rRNAs) and transfer RNAs (tRNAs), to perform target prediction analysis in the tomato genome. Among them, 7,042 *B. cinerea* sRNAs were predicted to target 3,185 distinct tomato mRNAs. Conversely, 114,011 tomato sRNAs were found that were predicted to target 11,434 *B. cinerea* mRNAs. Based on the sRNA-seq and mRNA-seq data, the authors observed a weak negative correlation between the entire set of sRNAs and their potential targets that was not significant according to a permutation analysis. Thus, they concluded that there was no evidence for cross-kingdom RNAi in the *B. cinerea-*tomato interaction. This conclusion is based on the assumptions that all sequenced *B. cinerea* sRNAs can contribute to cross-kingdom RNAi, that an important part of the plant gene expression changes observable by mRNA-seq are caused by fungal sRNAs, and that predictions of sRNA-mRNA interactions have low levels of false positives. We believe that these assumptions are not accurate for the following reasons:

First, not all sequenced sRNAs of *B. cinerea* are genuine extracellular sRNAs, not all extracellular sRNAs would translocate into the host cells, and not all sRNAs that enter host cells will form a functional silencing complex, e.g., with the host AGOs. Even within the same organism, not all sRNAs can silence their bona fide targets at any time. Many sRNAs, such as most mammalian miRNAs, can have an average of 300-400 predicted targets; some even have over 1000 predicted target genes. However, many studies have shown that only a small subset of these target genes’ transcripts are reduced in abundance at any given time under specific developmental stages or in response to various environmental cues. In contrast, most predicted target genes are not suppressed (Bartel, 2009, Wilczynska & Bushell, 2015). This phenomenon cannot lead to a conclusion that there is no evidence for mammalian miRNA to induce the silencing of target genes. Thus, conducting global sRNA-mRNA correlation analysis for cross-kingdom/cross-species RNAi would not yield meaningful results.

Second, the quality and reproducibility of Qin *et al*.’s sRNA- and mRNA-seq datasets are questionable. The sRNA data includes large amounts of reads mapping to rRNA and tRNA (up to over 90%) for all their analysis (see below for details). This could be an indication of low-quality and highly degraded RNA, and these sequences will mask any real silencing patterns. Restricting the analysis to sRNAs of 20-24nt, as per Qin *et al*.’s methods, will help increase the signal-to-noise ratio, as would excluding some reads mapping to rRNA/tRNA. Nevertheless, these steps will not effectively remove the problem, since all the RNA in these samples would have suffered similar levels of degradation, and even 20-24nt reads would include a large fraction of irrelevant degradation products.

Third, Qin *et al*. predicted tomato targets using the psRNATarget tool with the default settings of V2 2017, without any further requirements. These settings allow bulges and gaps in the target site base pairing and up to two mismatches in the seed region. An Expectation score of 5 was used as a cutoff. The stringency of these settings for plant target prediction is rather low because gene silencing rarely occurs with a bulge or gap in the sRNA-target pairing regions and mismatch at the 10^th^ - 11^th^ nt positions will block the function of sRNA (Schwab *et al*., 2005), which was the principal basis for designing the short tandem target mimic (STTM) to block the function of sRNAs (Todesco *et al*., 2010). To achieve more precise target prediction, in our previous paper (Weiberg et al., 2013), we used the TAPIR1.1 tool (Bonnet *et al*., 2010, Schwab et al., 2005) to predict tomato targets with more stringent requirements: a) No gap or bulge within the alignment between the sRNA and the target was allowed; b) The 10^th^ and 11^th^ nts of the sRNA must perfectly match its target; c) At most one mismatch or two wobbles were allowed from position 2 to 12; d) A maximum of two continuous mismatches was allowed; e) A score of 4.5 was used as a cutoff.

Thus, the extremely high number of tomato target genes predicted by Qin *et al*. (2022) was most likely overestimated. Furthermore, many of their predicted target genes in tomato may not be true Bc-sRNA targets. Importantly, even true sRNA/mRNA targets will not be functional unless at the very least, they are present in the same cell and within a silencing complex.

Most importantly, the essential step after the deep-sequencing and bioinformatics analyses is to perform experimental validation analysis, preferably using multiple approaches, to confirm all the conclusions drawn from any omics analysis. All the studies from our labs have followed this principle, and we always use molecular, biochemical, cell biological and/or genetics approaches to confirm any results we obtained from omics data.

### 2.2 Some *B. cinerea* sRNAs associate with tomato AGO1 supporting the presence of cross-kingdom RNAi between *B. cinerea* and tomato

We previously observed that a subset of *B. cinerea* sRNAs move into plant cells and utilize *Arabidopsis* AGO1 proteins to silence host target genes (Weiberg et al., 2013). Following *Arabidopsis* AGO1 co-immunoprecipitation and RNA isolation, the *B. cinerea* Bc-siR3.1, Bc-siR3.2 and Bc-siR5 were detected by a stem-loop RT-PCR method (Weiberg et al., 2013). The same molecular mechanism was also found in plant pathogens *F. oxysporum, V. dahliae, H. arabidopsidis*, animal fungal pathogen *Beauveria bassiana* and even in beneficial bacterial symbiont *Rhizobium* during cross-kingdom RNAi with their hosts (Ji et al., 2021, Zhang et al., 2022, Dunker et al., 2020, Ren et al., 2019). The sRNAs from these microbes are loaded into host AGO proteins to silence host target genes.

In order to experimentally validate whether cross-kingdom RNAi is present between *B. cinerea-*tomato during infection, the most straightforward way is to examine whether any *B. cinerea* sRNAs are associated with tomato AGO proteins after moving into tomato cells by performing tomato AGO1-immunoprecipitation (AGO1-IP) analysis. We utilized the tomato AGO1 antibody generated by the Katherine Borkovich lab, which was used to successfully pulldown tomato sRNAs (Ji et al., 2021), and immunoprecipitated the sRNAs associated with the tomato AGO1 after *B. cinerea* infection.

We indeed detected some incorporated Bc-sRNAs. The three Bc-sRNAs described in our previous study (Weiberg *et al*., 2013), Bc-siR3.1, Bc-siR 3.2, and Bc-siR5, were detected to be associated with tomato AGO1 complex at 12, 16 and 24 hpi (Figure 1a). We also checked Bc-sRNAs predicted to target tomato genes by Qin *et al*. (2022) and mapped them to annotated regions in the *B. cinerea* B05.10 genome. We found that two Bc-sRNAs (#3 and #5) were also associated with tomato AGO1 (Figure 1b). To determine whether these AGO1-associated Bc-sRNAs can trigger host target gene silencing, we examined the target gene expression after *B. cinerea* infection. We found that the tomato target gene of Bc-siR3.1, *Autophagy-related 2* (*SlATG2*); the target gene of Bc-siR3.2, *Mitogen-activated protein kinase kinase kinase* (*SlMPKKK4*); and the target gene of Bc-siR5, *Pentatricopeptide repeat protein* (*SlPPR*) was suppressed upon *B. cinerea* infection. Meanwhile, at least one of the Bc-siRNAs detected by Qin *et al*. (2022), the Bc-sRNA#3 tomato target, *Avr9/Cf-9–INDUCED F-BOX1* (*SlACIF1*), was down-regulated after infection (Figure 1c). Silencing of a Bc-sRNA#5 target, the *receptor-like kinases* (*SIRLK*) was also detected by Qin *et al*. in figure 4 of Qin *et al*. (2022). Most of these Bc-sRNA tomato target genes are associated with plant immune responses. For instance, ATG2 negatively affects powdery mildew resistance and mildew-induced cell death in Arabidopsis (Wang *et al*., 2011). Knockdown of *SlMPKKK4* using virus-induced gene silencing (VIGS) resulted in enhanced disease susceptibility in response to *B. cinerea* (Weiberg et al., 2013). The *SlACIF1* is involved in *Cladosporium fulvum* infection in tomato (van den Burg *et al*., 2008). Some PPR-derived siRNAs have been predicted to target genes in *Verticillium dahliae*, indicating their potential contribution to defense against fungal pathogen infection(Hudzik *et al*., 2020). Many of the RLKs have been found to detect microbe- and host-derived molecular patterns, serving as the first layer of defense responses (Tang *et al*., 2017). The transcript accumulation of their predicted tomato target genes was reduced during the infection, supporting that Bc-sRNA-mediated silencing of target genes occurred in tomatoes. Thus, we demonstrated that cross-kingdom RNAi is evident between *B. cinerea* and tomato. However, many Bc-sRNAs identified by Qin *et al*. (2022) were not associated with tomato AGO1 (#4, #6, #8, #9 in Figure 1b), which could explain why they did not silence the expression of corresponding predicted targets in tomato (Figure 1c).

**Figure 1.**
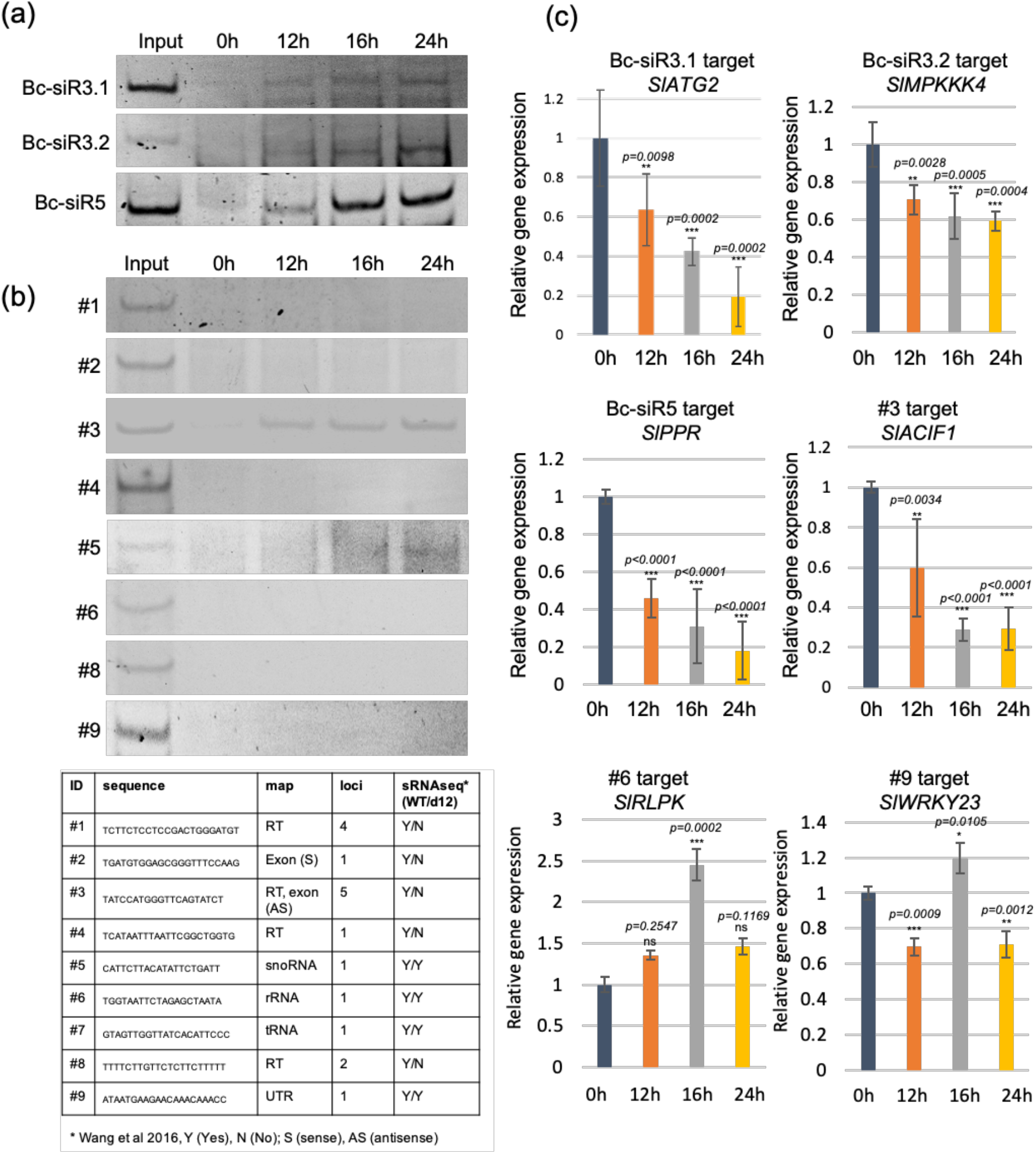
Detecting Bc-sRNAs by tomato AGO1 immunoprecipitation and tomato target gene expression during the infection. (a) and (b) Bc-sRNAs from our previous study (a) and from the Qin *et al*. (2022) study (b) were detected after tomato AGO1-IP, followed by RT-PCR. (c) Tomato target gene expression was detected using real-time PCR. RT, retrotransposons; UTR, untranslated region.

### 2.3 The sequencing data showed poor quality and a lack of reproducibility

To fully understand the results and conclusions from Qin *et al*. (2022), we decided to re-analyze their deep-sequencing data regarding quality and reproducibility. In order to avoid the appearance of potential bias, we invited another two independent hard-core bioinformatics groups (Jason Stajich and Cei Abreu-Goodger), both of whom have extensive experience in sRNA-seq analysis, to re-analyze Qin *et al*. ‘s sRNA-seq and mRNA-seq datasets. All three groups obtained similar results and reached the same conclusion: the overall reproducibility and the quality of Qin *et al*.’s deep sequencing data are low.

First, to assess the reproducibility and the quality of Qin *et al*.’s sRNA-seq dataset, we performed principal component analysis (PCA) and sRNA mapping distribution analysis. Only two or three libraries of the four biological replicates for each treatment were made available to the public, we included all the available libraries for the following analysis. The PCA analysis revealed no discernible differences between the samples and no conclusive clustering of replicates within the same treatment category. As shown in Figure 2a, the reproducibility among the biological repeats is low. For instance, one of the *B. cinerea* wild-type strain B5.10 cultured samples (B16) was grouped with one of the infected samples (I24), whereas another cultured *B. cinerea* sample formed a cluster group with both mock and infected samples. Qin *et al*. generated all the deletion mutants in the *B. cinerea ku70* mutant background (discussed below in details). Most surprisingly, *ku70*-3 was even grouped with *Δbcdcl1/Δbcdcl2/ku70-29* mutant, and *ku70-3* was very distinct from the other two mutant replicate libraries *ku70-1* and *ku70-2*. But *ku70-1* and *ku70-2* were clustered with the other two *Δbcdcl1/Δbcdcl2/ku70* mutants, which further indicates the low reproducibility of Qin *et al*.’s sRNA seq datasets. However, all samples with a *ku70* background, including *ku70* and the *Δbcdcl1/Δbcdcl2/ku70* mutants, are distinctly separated from all other samples using the wild type B05.10 *B. cinerea* strain. This result demonstrates that the sRNA profile of *ku70* differs significantly from B05.10. Figure 2b shows examples of the sRNA size and mapping distribution of the two biological repeats of cultured *B. cinerea* (upper panel, B16As and B16Bs), three biological repeats of the *ku70* mutants (middle panel *ku70-1, ku70-2, ku70-3*), and three biological repeats of the infected tomato sample collected at 12 hpi (lower panel I12As, I12Bs, I12Cs). The size and mapping distribution between the biological replicates are very different. For example, the first replicate of cultured *B. cinerea* (B16_1) shows a relatively uniform sRNA size distribution across 18-38 nt, with no obvious peak around 20-24nt. This is a typical pattern for degraded RNA, which is corroborated by the high levels of rRNA and tRNA-mapping reads. The first replicate of the *ku70* sample differed from the other two *ku70* replicates. In the first *ku70* replicate, a significant proportion of 22 nt sRNAs originated from TEs, while the other two *ku70* replicates showed a much lower proportion of sRNAs at this length. Additionally, the pattern of the *ku70* replicates was noticeably distinct from that of the cultured B05.10 *B. cinerea* samples, further demonstrating the big difference between the B05.10 and *ku70* mutant strains. The first two replicates of infected tomato samples (I12As and I12Bs), show a main peak at 33 nt, mainly consisting of tRNA fragments., while the third replicate (I12Cs) shows a substantially different profile (Figure 2b). If we assume that TE-derived sRNAs are likely to be responsible for cross-kingdom RNAi, the proportion of such reads varies drastically between replicates. We found that the vast majority of *B. cinerea* sRNAs claimed in Qin *et al*.’s datasets are derived from the sense strand of rRNAs and tRNAs (Figure 2c). For example, over 90% of the sRNAs from both cultured *Botrytis* libraries are from sense strands of rRNAs and tRNAs. Independently, we further analyzed the sRNAs mapped to the protein-coding regions (Figure 2d). Again, most of the sRNA reads are derived from the sense strand of the RNAs. This is another indication that these sRNAs detected are most likely degradation products.

**Figure 2.**
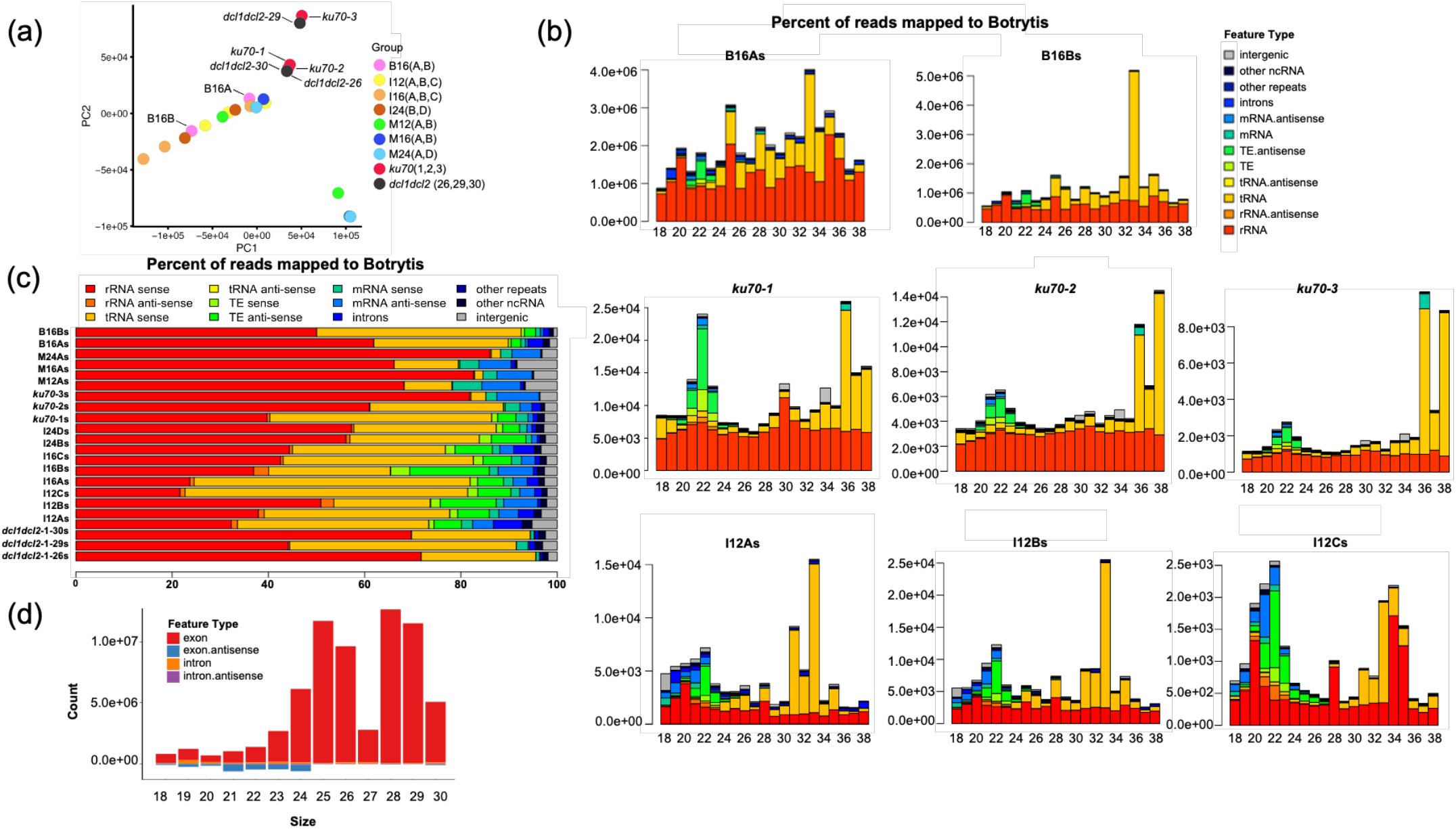
The reproducibility and quality of Qin *et al*. (2022) sRNA datasets are questionable. (a) Principal component analysis (PCA) of 22 *Botrytis* sRNA libraries. Please note that only 13 libraries were released in 2020, and the other 9 were released in 2023. Some treatment groups (M12, M16, M24, I24 and B16) only have two replicates, and the other groups (I12, I16, *Ku70, bcdcl1/bcdcl2*) have three replicates. Replicate names were denoted in the brackets after each treatment. (b) Genomic origin and size distribution of sRNAs from two *Botrytis* culture samples. Three infected tomato plant replicates and three *bcdcl1/bcdcl2* mutants show different patterns among biological replicates. (c) The distribution of reads from Qin *et al*. (2022) sRNA libraries that map to the different parts of the *Botrytis* genome. (d) The distribution of reads from Qin *et al*. (2022) sRNA libraries that map to coding regions. Coding of samples: I= inoculated tomato leaves; M=mock-treated tomato leaves; B= *B. cinerea* liquid culture; 12, 16, 24 = time point of sampling after inoculation (hours); A, B, C, D = biological replicate sample that was sequenced by Qin *et al*. (2022); s=sRNA fraction.

Second, we also performed PCA analysis to assess the reproducibility and the quality of the mRNA-seq dataset from Qin *et al*. (2022). Similarly, the biological repeats are not grouped according to the general biological expectation (Figure 3a). Mock libraries and *Botrytis-*infected libraries were grouped with each other (Figure 3a), indicating that the mRNA sequencing data have low repeatability and quality. To determine if the difference within each group is statistically significant, we performed differential expression analyses between infected and mock samples at each time point. We visualized results using volcano plots based on the FDR (false discovery rate) and foldchange value between different replicates and treatments. When using an FDR <0.05 and log2 fold change >1 as a cutoff, there are few (90 and 23) mRNAs with significant changes in “Infection 12 hours” and “Infection 24 hours,” respectively (Figure 3b-d). Thus, using these data to determine the expression levels of Bc-sRNAs targeted mRNAs is most likely inaccurate. The mRNA-seq datasets of *ku70* and the *Δbcdcl1/Δbcdcl2/ku70* were still not available, therefore, we were not able to check whether the silencing effect of sRNA targets of tomato was eliminated in the *dcl* double mutant.

**Figure 3.**
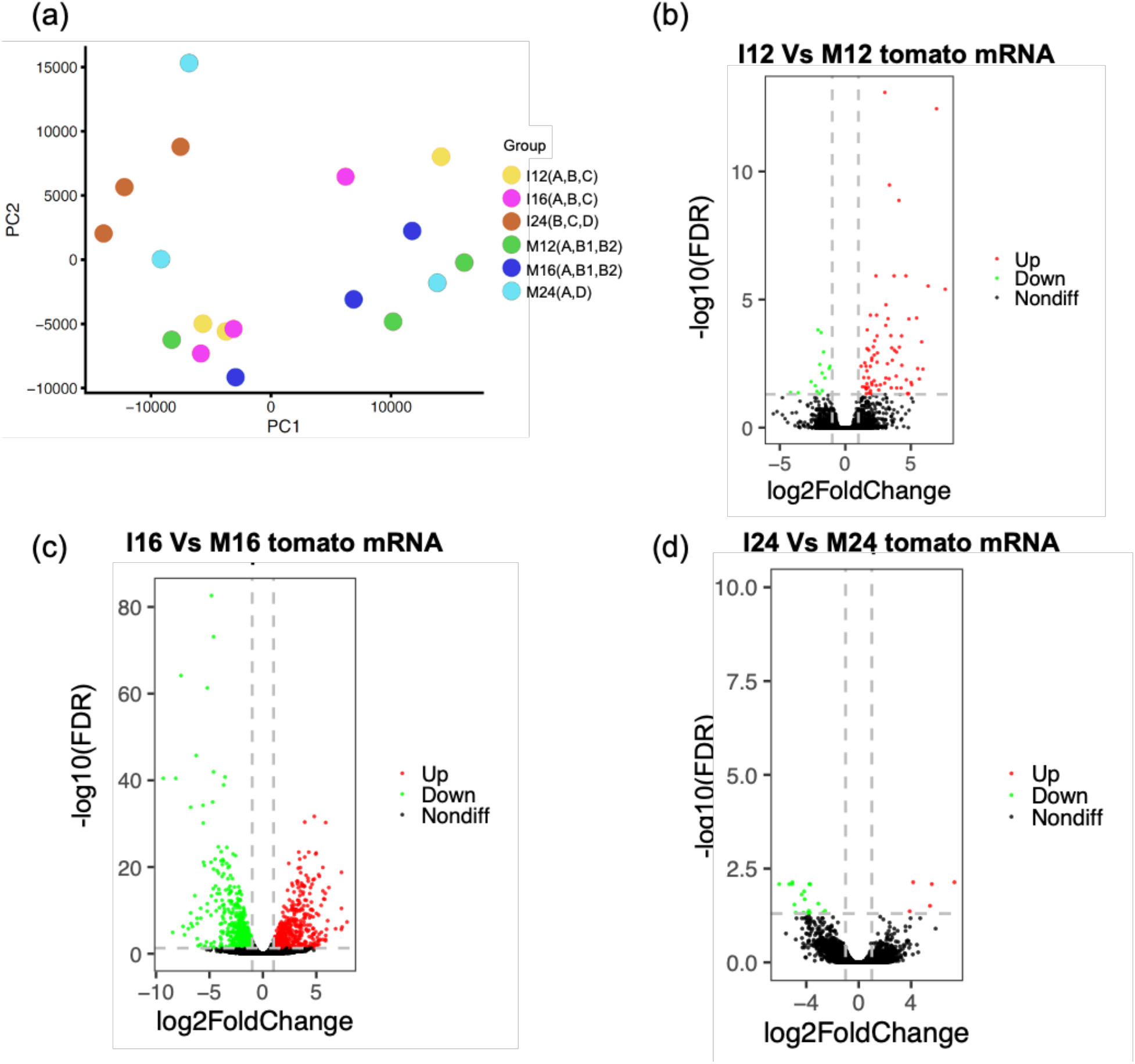
The reproducibility and quality of Qin *et al*. (2022) mRNA datasets are low. (a) Principal component analysis (PCA) of 18 tomato mRNA libraries. Please note that 15 libraries were released in 2020, and the other 3 were released in 2023. However, the 3 libraries (M12Bm, M16Bm and M24Dm) released in 2023 have the same name with 3 libraries released in 2020, so we renamed them as M12B1m, M16B1m and M24D1m (released in 2020), and M12B2m, M16B2m and M24D2m (released in 2023). (b-d) Volcano plot of differentially expressed genes (DEGs). The expression difference is significant for FDR <0.05 and |log2foldchange>1|. Coding of samples: I= inoculated tomato leaves; M=mock-treated tomato leaves; B= *B. cinerea* liquid culture; 12, 16, 24 = time point of sampling after inoculation (hours); A, B, C, D = biological replicate sample that was sequenced by Qin *et al*.; m= mRNA fraction.

### 2.4 sRNA production by the *ms3003* element

Qin *et al*. (2022) generated *ms3003* deletion mutant strains because “approximately 10% of the total sRNAs produced by *B. cinerea* strain B05.10 originated from a retrotransposon region on Chr14 annotated as *ms3003*”. Several questionable interpretations were made here:

First, estimating sRNA production per retrotransposon (RT) locus based on total sRNA read counts makes no sense because many sRNAs map multiple times and to different RT loci, leading to a doubling of sRNA read counts. Without knowing which element(s) is/are true sRNA-producing locus/i, the counts should be at least divided by the number of mapping events for estimation. Therefore, the 10% mentioned above of sRNA reads derived from the *ms3003* locus is misleading.

Second, there are even more fragmented RT elements (loci) - we have previously identified additional 221 RT loci of > 400 bp fragments (Porquier *et al*., 2021) dispersed over the *B. cinerea* genome, that have not been considered as a source of sRNA production by Qin *et al*., and which are making a valid estimation of the relative sRNA production per individual RT element/locus extremely complicated (if not impossible).

Third, it is not indicated by Qin *et al*. (2022), which sRNA library was used for estimating the percentage of sRNAs produced per RT element (as given in table.S3). This is highly relevant because we recently found that sRNAs mapping to different RTs, in particular belonging to the *Bc*Gypsy3 family, are highly up-regulated during tomato infection (Porquier et al., 2021).

There are contradictory results and interpretations in Qin *et al*. (2022). Deleting the *ms3003* element certainly did not lead to a drastic loss of sRNA production, as Qin *et al*. (2022) suggested. In fact, their data support the continuous production of RT-derived sRNAs, as shown in figure 5c of Qin *et al*. In order to make any statement about altered sRNA production, the authors should provide sRNA sequencing data for the *ms3003* mutants.

**Figure 4:**
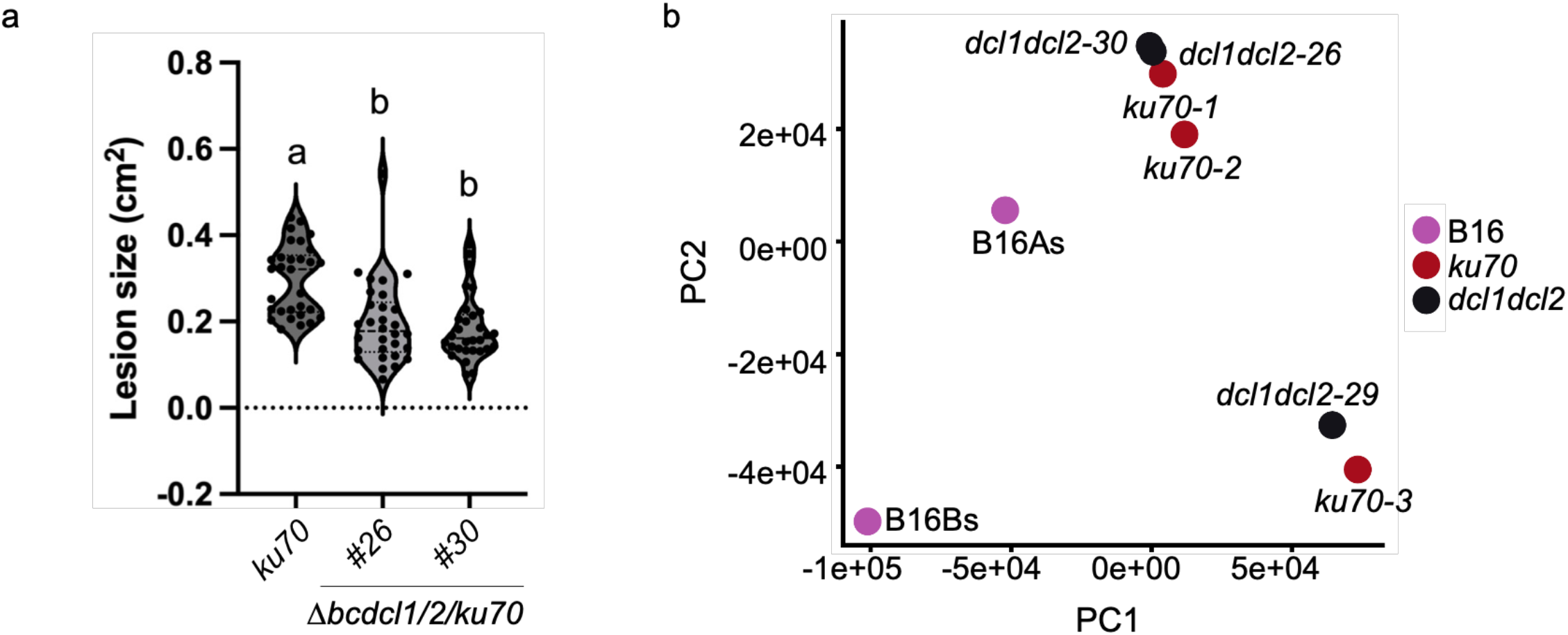
Infection assay of *Δbcdcl1/Δbcdcl2/ku70* on detached tomato leaves and PCA analysis of *ku70, dcl1dcl2*, and wild-type B05.10 strains. (a) Inoculation was performed with 100 conidiospores per inoculation, and lesion areas were analyzed at 60 hours post-inoculation (hpi). At least 30 lesions were measured per strain and condition, and inoculation assays were repeated three times with similar results. Letters indicate significant differences when using one-way ANOVA and Tukey test with p<0.05. (b) The small RNA sequencing libraries of the *ku70* mutant, *dcl1dcl2* mutant strain, and wild-type B05.10 strain were subjected to PCA analysis.

**Figure 5:**
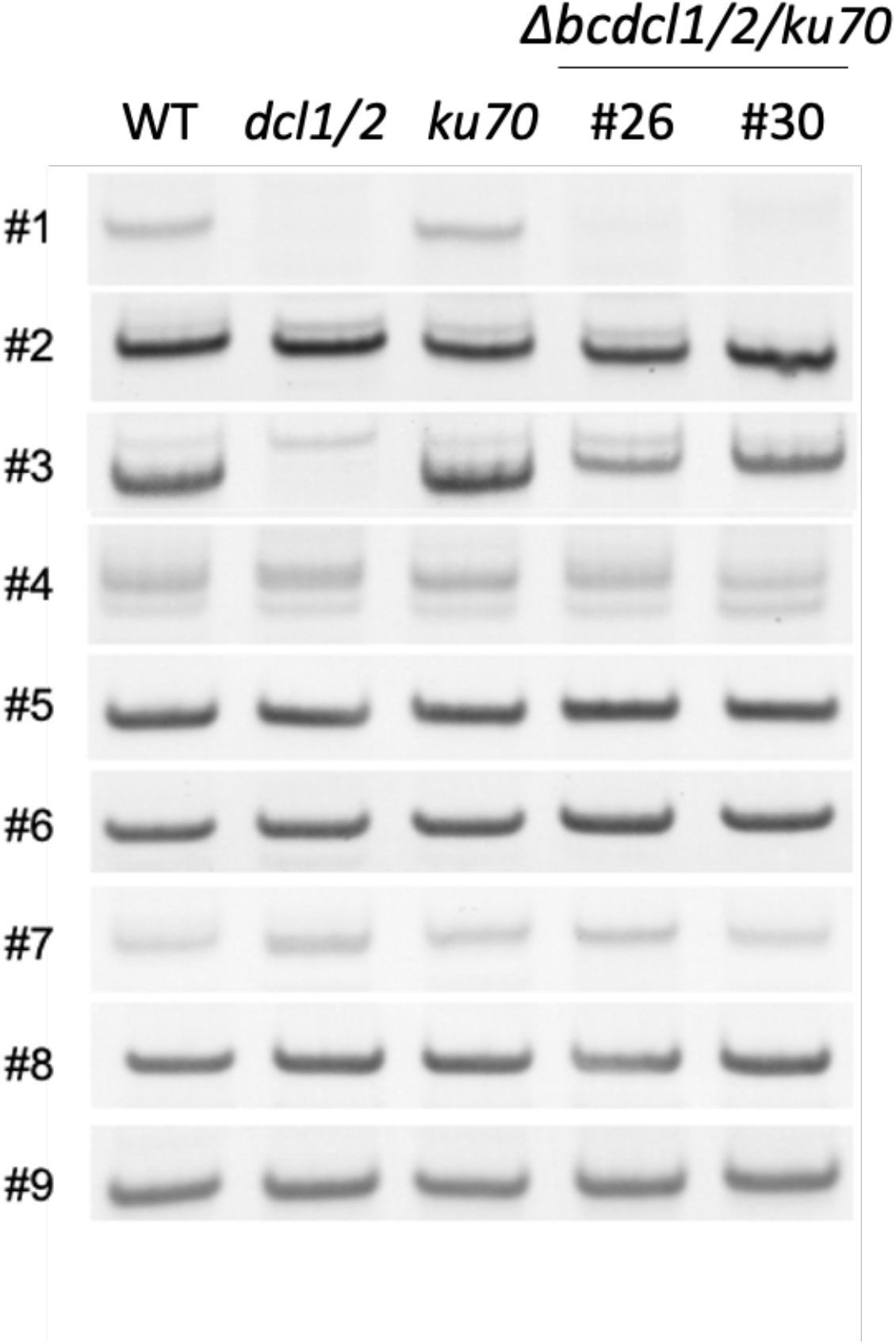
the *Δbcdcl1/Δbcdcl2* mutants produce sRNAs. Production of 9 BcsRNAs was tested in different *B. cinerea* genotypes by stem-loop RT-PCR. *dcl1/2* is the mutant strain generated by Weiberg *et al*. (2013), *Δbcdcl1/2/ku70* is the mutant strain generated by Qin *et al*. (2022).

### 2.5 Reproducibility of infection assay with the *Δbcdcl1/Δbcdcl2/ku70* mutant strains

Qin *et al*. argue that the newly generated *Δbcdcl1/Δbcdcl2* mutant in the *ku70* mutant background did not exhibit any reduction of virulence on tomato leaves and other tested plant species. For this experiment, they used a relatively high spore concentration of 1000 conidia/μl suspended in potato dextrose broth (PDB), a rich culture medium that allows rapid growth of *B. cinerea* in vitro, to inoculate the detached leaves. It is worth noting that tomato is a much more favored host for *B. cinerea* than Arabidopsis.

We have now repeated infection assays with detached tomato leaves using 10 μl of 10 conidia/μl spore concentrations, resuspended in 1% malt extract. We think this lower spore concentration better represents the natural infection pressure. Under this infection condition, we observed significantly reduced lesion formation in two independent *Δbcdcl1/Δbcdcl2/ku70* mutant strains at 60 hours post-inoculation (Figure 4A).

We would like to point out that the involvement of DCLs in fungal virulence was also observed in other fungal species and by other laboratories. Several studies reported that *dcl* deletion in fungal pathogens compromises infection, for example, in *Fusarium graminearum, Penicillium italicum, Colletotrichum gloeosporioides* and *Valsa mali* (Wang *et al*., 2017, Werner *et al*., 2021, Yin *et al*., 2020, Feng *et al*., 2017).

### 2.6 Qin *et al*. utilized an inappropriate NHEJ DNA repair mutant *ku70* as a background to generate *Δbcdcl1/Δbcdcl2* mutants

Qin *et al*. claimed that they generated *Δbcdcl1/Δbcdcl2* mutant strains using the Cas9 RNP strategy (Leisen *et al*., 2022) in a *B. cinerea* B05.10 *ku70* mutant background. This *B. cinerea ku70* mutant strain is compromised in a non-homologous end-joining DNA repair (NHEJ) system and favors homologous recombination to gain targeted gene knock-out (Choquer *et al*., 2008, Pinedo *et al*., 2008). However, double deletion of NHEJ and *DCL*s genes should be considered with caution for two reasons:

First, a *ku70* mutant may alter fungal pathogenicity. For example, the *ku70* mutant of the fungus *Penicillium digitatum* exhibited altered growth and conidia production (Gandía *et al*., 2016), which means that pleiotropic effects may occur due to increased sensitivity to DNA damage. In addition, RNAi mutants including *DCL*s showed increased sensitivity to DNA damage in *Neurospora crassa* (Lee *et al*., 2009), a phenomenon well-studied in the mammalian field as well (Tang & Ren, 2012, Bonath *et al*., 2018, Lu *et al*., 2018). Therefore, NHEJ plus *DCLs* deletion might likely cause unpredictable complications regarding DNA repair and genome integrity.

Indeed, PCA analysis showed that the sRNA profiles of the *ku70* mutant had huge differences from that in wild type *B. cinerea* B05.10 strain (Figure 4B).

Second, it might also be a problem when combining *ku70* mutant with CRISPR CAS9 system. CAS9 is known to generate unpredicted off-target cleavages, which largely rely on NHEJ DNA repair system. The combination of CAS9 and *ku70* mutation can generate more unpredicted mutation sites in the background. Many people in the field of mammalian and plant fungal pathogens are always hesitant in using *ku70/ku80* mutants because they are fully aware that knock-out NHEJ DNA repair function may cause many unpredicted mutations in the genome and also make the organisms more sensitive to DNA damage.

### 2.7 *B. cinerea Δbcdcl1/Δbcdcl2* still produces some transposon-derived *Bc* sRNAs

Qin *et al*. (2022) stated that “transposon-associated Bc-sRNAs are eliminated in the *Δbcdcl1/Δbcdcl2”*, according to their sRNA-seq data. We re-analyzed all the 22 sRNA-seq datasets, including cultured *B. cinerea*, the *ku70, Δbcdcl1/Δbcdcl2/ku70* mutant strains and *B. cinerea*-infected tomato. We found that sequencing depth among all libraries varies, with the *ku70* and *Δbcdcl1/Δbcdcl2/ku70* libraries displayed relatively low sequencing depths (Supplementary Table 1). We counted in all libraries reads of the nine Bc-sRNAs analyzed by Qin *et al*. in their figure.4 and figure.S1. 5 out of 9 tested Bc-sRNAs were neither detected in *ku70* nor *Δbcdcl1/Δbcdcl2/ku70* libraries, and the Bc-sRNA #6 was only detected in 1 out 3 *Δbcdcl1/Δbcdcl2/ku70* library replicates. The Bc-siRNA3.1, Bc-siR3.2, and Bc-siRNA5, as well as Bc-sRNA #5 and Bc-sRNA #6 revealed relatively high read counts compared to the other Bc-sRNAs (#1, #2, #3, #4, #7, #8, #9) in all *B. cinerea* libraries (Supplementary Table 1). Although accumulating in similar read counts, only Bc-siRNA3.1, BcsiR3.2 and Bc-siR5 bound to tomato AGO1, but not Bc-sRNA #6. Furthermore, the Bc-siRNA #3, which had relatively low read numbers, was still detected in tomato AGO1 (Figure 1b). These results highlight the importance of the tomato AGO1 co-IP sRNA analysis in order to identify potential cross-kingdom Bc-sRNAs.

Since the sequencing depth of *ku70* and *Δbcdcl1/Δbcdcl2/ku70* sRNA libraries were low, we designed PCR primers to detect the nine sRNAs, as analyzed by Qin *et al*. (2022), via stem-loop RT-PCR (Varkonyi-Gasic *et al*., 2007a). Herein, we detected all nine sRNAs in the B05.10 wild-type and *ku70* mutant strain and eight out of nine within the independent *Δbcdcl1/Δbcdcl2/ku70* strains (Figure 5). After mapping these nine Bc-sRNA candidates to annotated regions in the *B. cinerea* B05.10 genome (Figure 1b), we found only four Bc-sRNAs mapped to retrotransposons with one Bc-sRNA mapped to both a retrotransposon and an exon (Bcin16g04520) locus. Thus, many RT-derived sRNAs remain in the *Δbcdcl1/Δbcdcl2/ku70* mutants. Deleting *B. cinerea Bcdcl1* and *Bcdcl*2 cannot eliminate all transposon-derived sRNAs. Furthermore, our *Δbcdcl1/Δbcdcl2* double mutant differs from the *Δbcdcl1/Δbcdcl2/ku70* triple mutant that Qin *et al*. generated. Some *Botrytis* sRNAs, for example, Bc-sRNA#3 is still present in the *Δbcdcl1/Δbcdcl2/ku70* mutant but not in our *Δbcdcl1/Δbcdcl2* mutant (Figure 5). This Bc-sRNA can associate with tomato AGO1 protein to suppress the target gene expression, which strongly supports that these different *dcl* double mutants have different sRNA patterns and finally affect their virulence.

In summary, all the new experimental evidence presented here demonstrates that Qin *et al*.’s conclusion that cross-kingdom RNAi does not occur between *B. cinerea* and tomato is not supported by reliable experimental evidence or correct interpretation of existing data.

## 3 DISCUSSION

Small RNA-mediated RNA interference (RNAi) is a conserved eukaryotic gene-silencing mechanism. Accumulating evidence has shown that sRNAs can not only regulate gene expression in the same cells, but they can also travel between interacting organisms to induce gene silencing in trans, which is called cross-kingdom or cross-species RNAi (Halder *et al*., 2022, Wong-Bajracharya et al., 2022, Cui et al., 2019, Ren et al., 2019, Dunker et al., 2020, Shahid et al., 2018, Ji et al., 2021, Zhang et al., 2022, Buck et al., 2014, Chow et al., 2019, Sahr et al., 2022). Recently, Qin *et al*. (2022) stated that the sRNAs produced by the fungal pathogen *B. cinerea* might not contribute as much to cross-kingdom RNAi between *B. cinerea*-tomato interaction. We addressed the questions raised by Qin *et al*. (2022) and provided new experiments and analysis to prove that the conclusion reached by Qin *et al*. that cross-kingdom RNAi does not occur between *B. cinerea* and tomato is not supported by credible experimental evidence or accurate interpretation of the available data.

The basic theory of cross-kingdom or cross-species RNAi is that a specific group of sRNAs is selected and transported into the interacting organisms and bind to the RNAi machinery to then silence target gene expression. However, Qin *et al*. predicted the tomato target genes using all sRNAs identified in *B. cinerea* but ignored the sophisticated sRNA selection and delivery mechanism between *B. cinerea* and plants. This will cover up the actual effect of cross-kingdom RNAi-mediated gene suppression. Meanwhile, we experimentally validated the tomato AGO1 association with the sRNAs published in Qin *et al*.’s paper. We found that most of the sRNAs identified by Qin *et al*. are derived from rRNAs and tRNAs, and very few are associated with tomato AGO1, which can directly explain why those predicted sRNA target genes were not suppressed during *B. cinerea* infection.

Reanalysis of the raw sequencing data from Qin *et al*. by two independent bioinformatics groups found that the correlation efficiencies of the biological replicates and the quality of the datasets were inadequate to draw reliable conclusions. Most of the sRNA reads were derived from rRNA and tRNA regions or the sense strand of mRNAs, indicating that the sRNA detected by Qin *et al*. were degradation products.

The *Δbcdcl1/Δbcdcl2* generated by Qin *et al*. needs further careful characterization. A mutant strain, *ku70*, but not the wild-type *B. cinerea* strain was used to generate the mutant. This *ku70* mutant background can increase unpredicted background mutations when CRISPR-CAS9 mediated gene knock-out is applied. As demonstrated by the PCA analysis of the sRNA profile of *ku70* and the *Δbcdcl1/Δbcdcl2/ku70* mutants were very different from all B05.10 wild-type *B. cinerea* samples. The distribution of sRNA lengths in the *ku70* sample also differed from that of the cultured B05.10 *B. cinerea*, indicating a huge difference between these two strains. Additionally, our experimental validation has confirmed that not all the transposon-derived sRNAs have been eliminated from the Δ*bcdcl1/Δbcdcl2/ku70* triple mutant. Regarding the infection assay of *Δbcdcl1/Δbcdcl2/ku70*, more conditions should be considered, for example, different spore concentration and different host species for infection need to be taken into consideration. We observed significantly reduced lesion formation in two independent *Δbcdcl1/Δbcdcl2/ku70* mutant strains using lower spore concentrations for infection assay at 60 hours post-inoculation.

In summary, cross-kingdom RNAi is present between *B. cinerea* and tomato. Some *B. cinerea* sRNAs can travel into tomato cells and bind to tomato AGO1 to silence target genes in the host.

## 4 EXPERIMENTAL PROCEDURES

### 4.1 *Botrytis* and plant growth

The *B. cinerea* (Pers.: Fr.) strain B05.10, *ku70* mutant, and all three *Δbcdcl1/Δbcdcl2* mutant strains were cultured on HA medium (4 g/L yeast extract, 4 g/L glucose, 10 g/L malt extract 15 g/L agar) for leaf inoculation assay. The plates were incubated for 2 weeks at room temperature under the exposure of UV-Tube Complete Set, metal, 18W/60 cm (Eurolite^®^) to stimulate sporulation. Liquid-cultured *Botrytis* was used for RNA extraction and stem-loop PCR. 2×10^5^ conidia were cultured in 50 ml HA liquid medium at 22 °C for 20 h.

Tomato *S. lycopersicum* (L.) Heinz cultivar was used as a plant host in this study. Tomato seeds were incubated in a petri dish half-filled with sterile water in darkness at 22 °C for 3 days. Then the seeds were transferred into soil and grew for 5-6 weeks at 22 °C with 60% humidity and 16 h/8 h light/darkness.

### 4.2 Tomato AGO1 immunoprecipitation (IP)

Tomato AGO1 antibody from Dr. Borkovich’s lab was used to pull down the sRNAs associated with the AGO1 protein(Ji et al., 2021). Tomato AGO1 IP was performed with 5 g fresh leaves collected at 0, 12, 16 and 24 h after spray inoculation with *B. cinerea*. Tissues were collected and ground in liquid nitrogen before resuspension in 5 ml of extraction buffer (20 mM Tris–HCl at pH 7.5, 150 mM NaCl, 5mM MgCl_2_, 1 mM DTT, 0.5% NP40, proteinase inhibitor cocktail; Sigma). The extract was then precleared by incubation with 50 ul of Protein A-agarose beads (Roche) at 4°C for 30 min. The precleared extract was incubated with 10 ul of AGO1-specific antibody for 6 hours. A volume containing 50 ul of Protein A - agarose beads (Roche) was then added to the extract before overnight incubation. Immunoprecipitates were washed three times (5 min each) in wash buffer (20 mM Tris–HCl at pH 7.5, 150 mM NaCl, 5mM MgCl_2_, 1 mM DTT, 0.5% Triton x-100, proteinase inhibitor cocktail; Sigma). RNA was extracted from the bead-bound EVs with TRIzol reagent (Invitrogen). The sRNA RT–PCR was performed as previously described (Weiberg et al., 2013). Primer sequences are provided in Supplementary Table 2.

### 4.3 Bc-sRNA target gene expression analysis

The *B. cinerea* spores were diluted in 1% sabouraud maltose broth buffer to a final concentration of 1 × 10^5^ spores/ml. The diluted spores were evenly sprayed on 5- to 6-week-old tomato plants. The tomato leaves were collected after 0, 12, 16, and 24 hours of infection. Then total RNA was extracted from tomato leaf samples using TRIzol reagent (Invitrogen). The Bc-sRNA target gene expression was detected using quantitative real-time PCR. Primer sequences are provided in Supplementary Table 2.

### 4.4 Leaf inoculation

Tomato detached leaves were collected from 5-6 weeks old tomato plants. *B. cinerea* conidia were harvested from HA plate, where *Botrytis* had grown for 2 weeks. The conidia were adjusted the final concentration to 10 conidia/μl and resuspended in 1% malt extract for 1 hour. Tomato leaves were inoculated with 10 μl conidia suspension. Infected leaves were incubated in a humidity box. Infected phenotypes were photographed and quantified the lesion size using the Fiji software, ImageJ version 2.1.0/1.53c (Schindelin *et al*., 2012).

### 4.5 Reverse transcriptase PCR

CTAB-based RNA extraction method is used in this study (Bemm *et al*., 2016). Stem-loop PCR-based reverse transcription was performed with 1 μg total RNA using SuperScript™ III Reverse Transcriptase (Thermo Fisher Scientific) with small RNA specific primers (Varkonyi-Gasic *et al*., 2007b). Reverse transcription is performed with a thermos cycler and incubated at 16°C 30 min, followed by 60 cycles of 30°C 30 sec, 42°C 20 sec, 50°C 1 sec. Samples were in the end heated up at 85°C 15 sec then kept at 4 °C for short storage. Stem-loop PCR was performed using GoTaq G2 Polymerase (Promega) together with stem-loop universal reverse primer and sRNA-specific forward primer.

### 4.6 RNA seq data analysis

#### 4.6.1 RNA seq data analysis method 1-Cei Abreu-Goodger

##### Processing of sRNA-Seq reads

A total of 22 sRNA-Seq raw sequencing files were downloaded from the NCBI Sequence Read Archive under BioProject PRJNA496584. Files in fastq format were extracted using fastq-dump from sra-tools (NCBI, 2022). Sequencing quality was analysed using FastQC (Andrews, 2010) and MultiQC (Ewels *et al*., 2016), and was judged to be of good quality for all samples. The NEBNext 3’ adapter was trimmed using reaper (‘-geom no-bc -clean-length 18 -3p-global 12/2/1 -3p-prefix 8/2/1 -3p-head-to-tail 1 -format-clean >%I%n%C%n -nnn-check 1/1 -qqq-check 10/1/0/0 -3pa AGATCGGAAGAGCACACGTCT) (Davis *et al*., 2013). Trimmed reads between 18-38 nt were collapsed to unique sequences using tally while preserving their counts from each sample (-l 18 -u 38 -format ‘>seq%I_w%L_x%C%n%R%n’) (Davis et al., 2013). Sequences that were only observed 1 or 2 times across the 22 samples were discarded as they most likely represent sequencing errors, and in any case contain little useful information.

##### Mapping and assigning to genome of origin

Genome sequences for the *Botrytis cinerea* B05.10 (ASM83294v1) and *Solanum lycopersicum* SL3.0 where downloaded from Ensembl v55 (Yates *et al*., 2022), and unique sequences were mapped using bowtie1 (Langmead *et al*., 2009), allowing 0 or 1 mismatch, saving those that mapped or not to each genome but without further mapping information. Sequences were assigned to the genome with fewer mismatches, to “both” if the number of mismatches was the same and “none” if no hit was found. All sequences assigned only to *B. cinerea* were mapped again with bowtie1 using the parameters -v 1 -k 10000 --best --strata. The read counts assigned to each sequence in each sample were divided equally among all mapping locations.

##### Botrytis genome annotation

Genomic coordinates for protein-coding genes, tRNA, rRNA, other non-coding RNA classes and pseudogenes, were taken from the Ensembl GFF file. Transposable Element annotation was obtained by running RepeatModeler2 (Flynn *et al*., 2020) and RepeatMasker (Smit *et al*., 2013-2015). To help assigning mapped sequences to unique annotations (and avoid overcounting), a non-redundant genomic annotation was built where all regions with overlapping annotations were reduced to a single annotation. The following preference was used: rRNA > tRNA > Transposable Elements > protein-coding exons > other ncRNA > introns > pseudogenes. The counts for all sequences in a sample (Figure 2c), or divided by length (Figure 2b), were assigned to the annotation category at the central base of each mapping location. Data processing and plotting was done in R(Team., 2022), making use of Biostrings (H. Pagès, 2021), GenomicRanges(Lawrence *et al*., 2013), Rsamtools(Morgan M, 2021) and rtracklayer(Lawrence *et al*., 2009).

#### 4.6.2 RNA seq data analysis method 2- Jason Stajich

Bioinformatic analyses to examine read size distribution and genomic context were performed with scripts developed and available in this github repository https://github.com/stajichlab/Botrytis_sRNA_profile. Sequence reads were processed with fastp (Chen *et al*., 2018) to detect and trim adaptors and remove low quality reads. Trimmed sequences were aligned to *Solanum lycopersicum* ITAG4 (doi: 10.1101/767764v1) downloaded from SolNet solgenomics.net and *Botrytis cinerea* B05-10 genome downloaded from FungiDB (Basenko *et al*., 2018) using STAR aligner (Dobin *et al*., 2013). The resulting SAM alignments were converted to BAM files with samtools and processed with a custom python scripts ‘process_BAM_sRNA.py’ and ‘get_BAM_sRNA_unique.py’ which relied on Biopython (Cock *et al*., 2009) and pybedtools (Dale *et al*., 2011) generate size distribution and classifications of genomic features overlapped by the read alignments and visualized as plots with custom R scripts ‘plot_sRNA_sizes.R’ and ‘plot_sRNA_sizes_uniq.R’.

#### 4.6.3 RNA seq data analysis method 3- Qiang Cai

Data was acquired from the NCBI database (https://www.ncbi.nlm.nih.gov/) with accession number PRJNA496584. The data was detected quality by using FastQC v0.11.9. Adapter clipping, low-quality bases removal and reads size selection were applied to the data with Trim Galore v0.6.10. The clean data was used to align to the *B. cinerea* B05.10 genome ASM83294v1 assembly and *S. lycopersicum* genome SL3.0 assembly by using bowie v1.3.1. The reads unmapped to *S. lycopersicum* genome and exactly mapped to the *B. cinerea* B05.10 genome were used for subsequent analysis. After counting sRNA reads, the CPM (Counts Per Million mapped reads) was calculated by using R package edgeR v3.36.0. Principal component analysis (PCA) was applied for the above samples with the R package PCAtools v2.6.0 to obtain more insight into the separation between samples of each treatment separately. The ggplot2 v 3.4.0 package was used to draw a relative image in R.

##### *S. lycopersicum* transcriptome analysis

Data was acquired from the NCBI database (https://www.ncbi.nlm.nih.gov/) with accession number PRJNA496584. The above sequencing reads were filtered using fastp v0.23.2 with adapter trimming and quality filtering function. The filtered reads were mapped to *B. cinerea* genome ASM83294v1 assembly and *S. lycopersicum* genome SL3.0 assembly using hisat2 v2.2.1. Then the reads unmapped to the *B. cinerea* genome and mapped to the *S. lycopersicum* genome were counted using featureCounts v2.0.1 and quantified in R. The original statistical expression matrix was normalized by RPKM (Reads Per Kilobases of exon model per Million mapped reads).

Principal component analysis (PCA) analysis was applied with the R package PCAtools v2.6.0 to reflect the repeatability between all samples. DESeq2 v1.34.0 was used to differentially express gene analysis and ggplot2 v3.4.0 was applied to draw the volcano plot. Through the above method, the differentially expressed genes which met the condition that the absolute value of log2-fold change was more than 1 and the FDR value was less than 0.05 were identified.

## Competing interests

The authors declare no competing interests.

## Acknowledgments

We thank Dr. Jan van Kan for providing us with the *B. cinerea* mutant strains *ku70* and the *Δbcdcl1/Δbcdcl2* #26 and #30. Work in the Q.C. laboratory was supported by grants from National Natural Science Foundation of China (32272029), Hubei Provincial Natural Science Foundation of China (2022CFA079). Work in the H.J. laboratory was supported by National Institute of Health (R35GM136379), National Science Foundation (IOS 2020731), United State Department of Agriculture (2021-67013-34258), United States Department of Agriculture National Institute of Food and Agriculture (2019-70016-29067) and the CIFAR ‘Fungal Kingdom’ fellowship.

## Data Availability Statement

All the data generated from this study will be made available to the public.

## Notes

### Competing Interest Statement

The authors have declared no competing interest.

### Summary of Updates

One more reference was added.

